# Collective behaviour is not robust to disturbance, yet parent and offspring colonies resemble each other in social spiders

**DOI:** 10.1101/761338

**Authors:** David N. Fisher, James L.L. Lichtenstein, Raul Costa-Pereira, Justin Yeager, Jonathan N. Pruitt

## Abstract

Groups of animals possess phenotypes such as collective behaviour, which may determine the fitness of group members. However, the stability and robustness to perturbations of collective phenotypes in natural conditions is not established. Furthermore, whether group phenotypes are transmitted from parent to offspring groups is required for understanding how selection on group phenotypes contributes to evolution, but parent-offspring resemblance at the group level is rarely estimated. We evaluated robustness to perturbation and parent-offspring resemblance of collective foraging aggressiveness in colonies of the social spider *Anelosimus eximius*. Among-colony differences in foraging aggressiveness were consistent over time but changed if the colony was perturbed through the removal of individuals, or via their removal and subsequent return. Offspring and parent colony behaviour were correlated, but only once the offspring colony had settled after being translocated. The parent-offspring resemblance was not driven by a shared elevation but could be due to other environmental factors. Laboratory collective behaviour was not correlated with behaviour in the field. Colony aggression seems sensitive to initial conditions and easily perturbed between behavioural states. Despite this sensitivity, offspring colonies have collective behaviour that resembles that of their parent colony, provided they are given enough time to settle into the environment.

## Introduction

Many organisms form groups (Ward and Webster 2016). These aggregations can help individuals avoid predation, acquire resources, find mates, and so on (Bilde et al. 2007; Frank 2007; Dobson et al. 2012; Almberg et al. 2015; Groenewoud et al. 2016). For many of these purposes, groups use collective behaviour, where individuals act in a co-ordinated or synchronised manner (Sumpter 2006). Collective behaviours cannot always be understood in terms of a simple sum of the actions of individuals and so groups can possess phenotypes that simply do not exist at the individual level (Parrish and Edelstein-Keshet 1999; Modlmeier et al. 2014; Farine et al. 2017). Group phenotypes are therefore a tier of biological organisation that require direct study, both in terms of how they relate to selection and evolution at the individual level, as well as in and of themselves (Couzin 2009).

Individual traits can range from being highly consistent within an individual to highly variable (Bell et al. 2009). An individual might retain its behaviour in spite of a disturbance, or it might find its behaviour changed as a result of a disturbance (Tuomainen and Candolin 2010; Sih et al. 2011). The same could be true of group phenotypes; the collective behaviour of groups may resist disturbances, or it may be altered by them (Flack et al. 2005, 2006; Smith et al. 2013; Kubitza et al. 2015; Formica et al. 2016). For instance, collective behaviours might be “self-organised”, where individuals re-create the same group behaviour after disturbances by following the same set of interaction patterns that created the initial group behaviour (Bonabeau et al. 1997; Fisher and Pruitt 2019; Fisher et al. 2019). In contrast, groups might change their behaviour following disturbances, if they are shunted into different “states” following a disturbance (Flack et al. 2005, 2006; Doering et al. 2018; Pruitt et al. 2018), or engage in non-linear interactions that give divergent trajectories, and so different group phenotypes, from a similar set of starting conditions (May and Oster 1976; Cole 1994; Fisher et al. 2018; Honegger and de Bivort 2018). However, the robustness of group phenotypes to disturbances is not well documented (Flack et al. 2005, 2006; Smith et al. 2013; Kubitza et al. 2015; but see: Formica et al. 2016).

If group phenotypes are resistant to disturbances and stable over time, then they can influence the survival and reproductive success of individuals within those groups (Wray et al. 2011; Keiser and Pruitt 2014; Pruitt and Goodnight 2014; Pruitt et al. 2017, 2019). Stability in group phenotypes is important because it determines the degree to which they can be subject to natural selection (in a population of groups, if all group phenotypes vary widely these phenotypes cannot be associated with relative fitness). One of the most extreme forms of group disturbance is group fission, whereby a subset of group members disperse or bud off to form a smaller, “daughter” group (Vollrath 1982; Aviles 1986). The collective behaviour of these daughter groups can be similar to that of their parent group and so exhibit a crude kind of collective or group-level heritability (Bienefeld and Pirchner 1990; Pruitt et al. 2017, 2019). However, unlike individual-level traits (Houle 1992), the heritability of group-level traits is not widely documented. This therefore makes it hard to judge how, if at all, group-level selection can contribute to evolution and adaptation (Wilson 1997b,a; Gardner and Grafen 2009; Queller and Strassmann 2009).

We therefore had two questions surrounding collective behaviour. First, is collective behaviour robust to disturbance? Second, is collective behaviour transmitted from parent group to offspring group in staged fission events? If both of these are true, then we might expect group phenotypes such as collective behaviours to play a more important role in evolution than is currently thought. We investigated these questions in a Neo-tropical social spider, *Anelosimus eximius* (Araneae: Theridiidae). *Anelosimus eximius* is classified as “non-territorial permanent social” (Avilés 1997), where individuals (sometimes numbering into the 10,000s; Avilés 1997) from overlapping generations live together in the same web structure and cooperate in web-building, prey capture, and alloparental care (Vollrath 1986; Ebert 1988; Avilés and Tufiño 1998; Avilés and Harwood 2012; Avilés and Guevara 2017; Pruitt and Avilés 2017). This allows them to feed on larger prey than would be expected of a spider of their body size and to endure environments where related species with lower levels of sociality cannot (Guevara and Avilés 2015; Avilés and Guevara 2017; Fernandez-Fournier et al. 2018). Once prey make contact with the web, social spiders collectively rush to immobilise it. How quickly the colony responds to a potential prey item can be an important determinant of colony success and so this is the collective behaviour that we focus on here (hereafter “foraging aggressiveness”; Lichtenstein et al. 2019).

## Methods

### Data collection

Our study took place in June and July 2019, near Tena, Ecuador (Fig. 1), under the Ecuadorian Ministry of the Environment permit no. 014-2019-IC-FLO-DNB/MA. We located colonies of *A. eximius* on roadsides, where they are relatively conspicuous on hedgerows, fences, and in trees. Their webs are composed of a “basket” at the base, with a sheet and tangle capture web above (Yip et al. 2008). Once we found colonies, we marked their location and recorded GPS coordinates to allow us to re-locate them. We then recorded their elevation and measured the height, width and depth of the basket. We found 45 colonies that were suitable for our study, being within reach of an observer and located within a morning’s drive of our laboratory. We tested these 45 colonies’ foraging aggressiveness three times over six days (every other day). Our test for foraging aggressiveness was the colony’s speed to attack a vibrating stimulus (following: Lichtenstein et al. 2019). We stimulated colonies to attack by touching a piece of wire fixed to a modified handheld vibratory device (8” Vibrating Jelly Dong, Top Cat Toys, Chatsworth CA, USA) to a small piece of leaf placed in the web. The leaf was always placed on the edge of the basket of the web, and we waited at least 60 seconds from the placement of the leaf before introducing the vibrations. The vibrations running through the leaf simulate a prey item caught in the web; assays similar to this are often used to estimate foraging aggressiveness in social (e.g. Laskowski and Pruitt 2014; Lichtenstein et al. 2019) and solitary (Dirienzo and Montiglio 2016; Montiglio and DiRienzo 2016) spiders. We timed the number of seconds from the start of the vibrations until a spider touched the leaf. If the colony did not respond within 10 minutes the score was set at 600 (2.3% of all trials). This test is repeatable among-colonies over four days (r = 0.26) and, at high altitudes, influences colony survival over a 11 month period (Lichtenstein et al. 2019), indicating it captures relatively stable aspects of colony collective behaviour.

**Figure 1.**
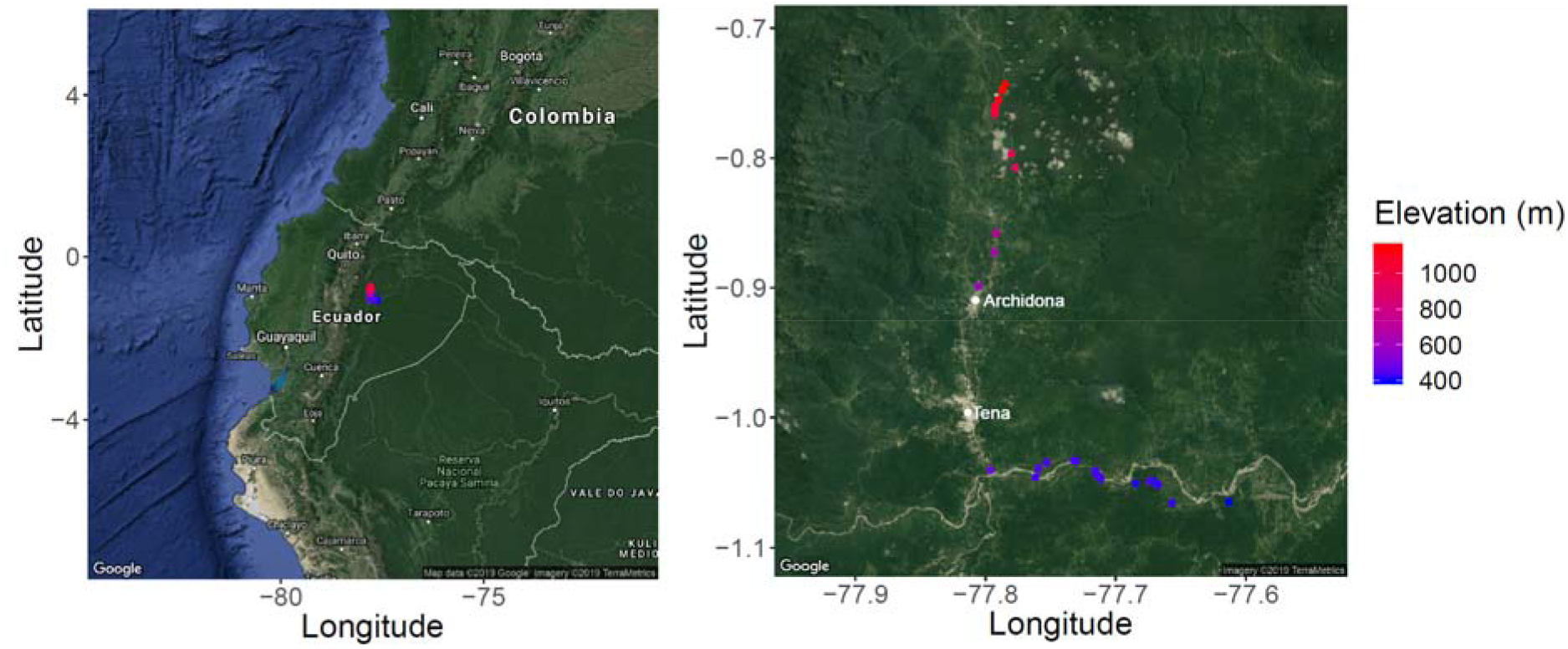
Maps showing the location of each of the *Anelosimus exemius* colonies in the study, with the elevation of the colony indicated by the colour (red = high elevation, blue = low elevation). In the right map the towns of Tena and Archidona are indicated with white points.

After these three baseline collective aggressiveness tests, we assigned each colony randomly to one of three treatments. Fifteen colonies were “removal”, 15 “procedural control” and 15 “control”. For the removal and procedural control colonies, we returned three days after the 3^rd^ behavioural test and removed a subset of spiders from each colony, placed them in sealed plastic boxes (190 x 190 x 90 mm) with sticks to support web building, and transported them back to our laboratory. Individuals were collected either by gently shaking the web and caching spiders that dropped or scooping a small bit of webbing into a large plastic box. We counted the number of individuals that were large (>2mm in body length), medium sized (<2mm & >1mm in body length) or small (<1mm in body length), with size being estimated by eye. We endeavoured not to destroy any vegetation the web was built on, in order to preserve the web’s structure. Control colonies were left undisturbed.

Each subset of spiders that we collected was left undisturbed to acclimatise to captivity in their box for two days. Boxes had four airholes to provide oxygen, and spiders were provided a moist piece of paper on the 4^th^ day of their captivity for hydration; they were not fed. We then tested the foraging aggressiveness of each of the 30 captive colonies three times over six days (every other day; the 1^st^ laboratory test beginning five days after the last pre-disturbance test). We modified the assay slightly to account for the new setting: we reduced the power of the vibrations to avoid over-amplification in the small box, and the wire was touched directly to the web rather than to a small leaf. These laboratory assays were used to assess the resemblance of parent and daughter colonies in a common garden environment. Although we might expect behaviour in the laboratory to differ substantially from that in the field, due to the lack of all natural cues (but see: Boon et al. 2008; Herborn et al. 2010; Fisher et al. 2015; Yuen et al. 2016), we might still expect the ranking of colonies in terms of their foraging aggression to be similar in both the laboratory and in the field. In this case a positive correlation would be expected.

Following their 3^rd^ test (on the same day), the spiders from procedural control colonies were placed directly back into their source (parental) colony. The colonies in this treatment group therefore lost no spiders but experienced the physical disturbance of the sampling event. Spiders from the removal treatment were placed in vegetation similar to what the parent colony had built its web on, but 5-10m away from the parent colony. This was designed to mimic the fission of a colony and the foundation of a new colony by a subset of individuals (sociotomy), which occurs naturally in *A. eximius* as colonies grow in size (Vollrath 1982; Venticinque et al. 1993; Avilés 1997). These “bud colonies” were used to assess the heritability of colony behaviours when in the same environment as their parent colony. At this point we discovered that eight of the parent colonies had been destroyed by workers clearing roadsides. Two of these colonies were in the procedural control group, but we could not return the previously removed spiders to a now destroyed colony, so we placed these spiders into vegetation 5-10m away as bud colonies.

Two days after returning them to the wild, we tested the collective aggressiveness of each surviving parent colony (n = 37) and each bud colony three times over six days (every other day) using the same method as before. In three instances the bud colony was completely abandoned, leaving 14 bud colonies (including the additional two colonies that were originally part of the procedural control group) to assay for foraging aggressiveness. To evaluate the robustness of *A. eximius* colonies to disturbance, we tested for a correlation between parent colonies’ pre- and post-disturbance behaviours. We evaluated transmission of aggressiveness from parent to daughter group by testing for a correlation between the pre-disturbance behaviour of parent colonies and the behaviour of bud colonies in a common garden setting (the laboratory) and a natural setting (the bud colony behaviours). During the three tests of the bud colony foraging aggressiveness, we observed the bud colonies frequently changing position and orientation in the vegetation. We thought it was likely that there was an initial “settling” phase after returning the bud colonies to the wild from captivity. Therefore, starting eight days after their 3^rd^ test, we tested each bud colony another three times over six days (every other day). This procedure was meant to capture bud colony behaviour following a settlement period (“settled bud behaviour”, the initial three tests hereafter being referred to as “initial bud behaviour”). A schematic outlining the sampling regime for the study is shown in Fig. 2.

**Figure 2.**
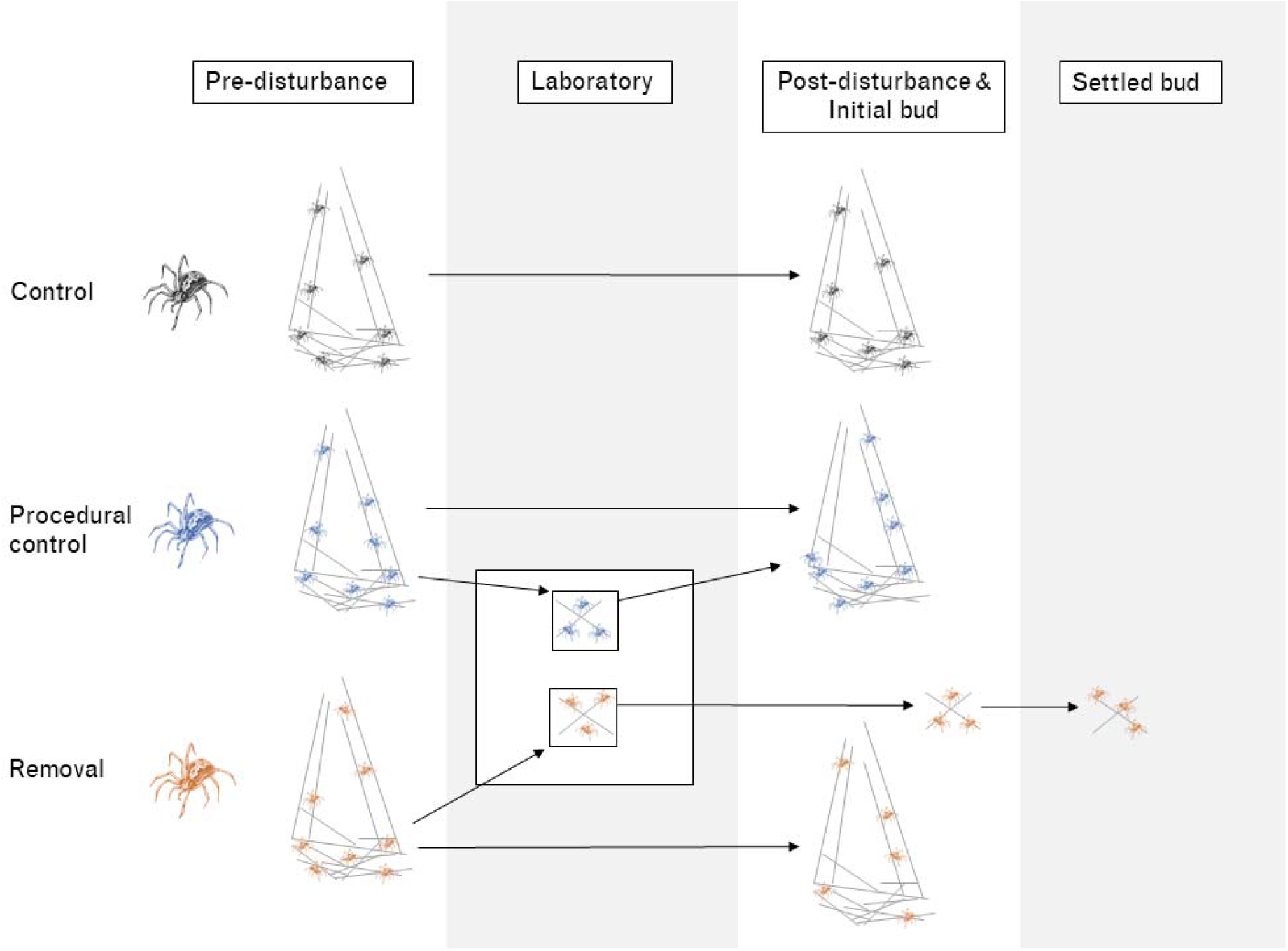
A schematic demonstrating our study design. In the pre-disturbance phase 45 colonies were tested three times over six days for foraging aggressiveness. For two thirds of these colonies (in the “removal” and “procedural control” groups) spiders were then removed to the laboratory, where they were tested three times over six days for foraging aggressiveness. Following this, spiders in the procedural control groups were returned to their original colony, while spiders in the removal groups were placed near the original colony as “bud colonies”. We then tested all original colonies and all bud colonies three times over six days. Following this we tested each bud colony another three times over six days to measure “settled” behaviour.

### Data analysis

To assess the stability of colony behaviour over time in face of the disturbance, we initially estimated the phenotypic correlation (Pearson’s correlations in all cases) between the log of pre-disturbance foraging aggressiveness and the log of post-disturbance foraging aggressiveness, with a colony’s first measure pre-disturbance paired with its first measured post-disturbance, and so on. However, this does not estimate the among-colony correlation between pre- and post-disturbance behaviours, instead it conflates among-colony, among-date and residual variation (analogous to the “individual gambit”; Brommer 2013; Dingemanse and Dochtermann 2013). To directly estimate the among-colony correlation between pre- and post-disturbance foraging aggressiveness, we built multivariate models in MCMCglmm (Hadfield 2010) with the logs of pre-disturbance foraging aggressiveness and post-disturbance foraging aggressiveness as response variables. We entered “NA” for the post-disturbance trials for colonies that had been destroyed. This allowed us to include their scores for the pre-disturbance trials in the model, which should improve the estimate of the among-colony variance in pre-disturbance foraging aggressiveness. We estimated the among-colony variances and covariance between pre- and post-disturbance foraging aggressiveness, the among-date variances for these traits (but no covariance as the two behaviours were never tested on the same day) and the residual variances for each behaviour (but no covariance as the two behaviours were never measured at the same time). We included the log of colony basket volume (height*depth*width), mean centred and scaled to a variance of one, and the trial number (1-3), mean centred, as fixed effects for each behaviour. This was done in case colony foraging aggressiveness covaried with size (Yip et al. 2008; Pruitt et al. 2011) and in case the colonies changed their behaviour over time.

To test if the disturbed colonies changed their behaviour more than the control colonies, we estimated the raw phenotypic correlations for each of the three treatment groups. We then we fitted the multivariate model described above to each of the three treatment groups separately and compared the magnitude and distributions of the among-colony correlations. If the control group had a stronger correlation between pre- and post-disturbance foraging aggressiveness than the removal or the procedural control groups, we could conclude that the disturbance disrupted colony collective behaviour.

To assess the resemblance of collective behaviour between parent and offspring colonies, we first estimated the phenotypic correlations between log-transformed pre-disturbance foraging aggressiveness, log-transformed laboratory foraging aggressiveness, and log-transformed bud colony foraging aggressiveness, associating the first pre-disturbance trial, the first laboratory trial, and the first bud trial and so on. However, phenotypic correlations such as this (including those based on only a single measure of parents and offspring, e.g. Pruitt et al. 2017, or those based on averages of parent and offspring colony traits, e.g. Pruitt et al., 2019) conflate among- and within-colony covariance, when only the former is relevant for assessing whether more aggressive parent colonies have more aggressive daughter colonies (Brommer 2013; Dingemanse and Dochtermann 2013; see also Niemela and Dingemanse 2018 for a discussion of the issues with using a single measure of behaviour to estimate covariances). To estimate the among-colony correlation, we built multivariate models in MCMCglmm, with the logs of pre-disturbance foraging aggressiveness, laboratory foraging aggressiveness, and bud foraging aggressiveness as response variables. We estimated the among-colony variances and covariance between these three traits. This is analogous to a parent-offspring regression, which overestimates heritability compared to estimates from an “animal model” (Kruuk 2004). We did not have a colony level pedigree, nor could we calculate the relatedness among colonies by some other means. Therefore, the parent-offspring covariance we estimate here should be taken as an upper limit for the true colony level heritability.

We also estimated among-date variance for each behaviour (but no covariance as the behaviours were never tested on the same day) and the residual variance for each behaviour (but no covariance as the behaviours were never measured at the same time). We included the log of colony volume as a fixed effect for pre-disturbance behaviour, and the number of adults removed from the colony and so tested in both the laboratory and as a bud colony (summing large and medium spiders, so any greater than 1mm in body length) as fixed effects for laboratory and bud behaviour. This was done in case colony size impacted foraging aggressiveness. These fixed effects were scaled to a mean of zero and a variance of one. We also include trial number (1-3) as a fixed effect, mean centred, in case the colonies changed their behaviour over time.

We estimated the raw phenotypic correlations once with the 1^st^-3^rd^ tests on the bud colonies (initial bud behaviour) and once with the 4^th^-6^th^ tests (settled bud behaviour). We also re-fitted the multivariate model using the 4^th^-6^th^ tests instead of the 1^st^-3^rd^ tests. If collective behaviour was inherited from parent colony to offspring colony, we expected a positive among-colony correlation between the pre-disturbance and bud behaviours. If behaviour in the laboratory reflects behaviour in the field, then there would also be a positive among-colony correlation between the pre-disturbance and laboratory foraging aggressiveness. Further, if the 4^th^-6^th^ tests on the bud colonies reflects settled behaviour, we expected the among-colony correlation between pre-disturbance foraging aggressiveness and the settled bud foraging aggressiveness to be stronger than the correlation between pre-disturbance foraging aggressiveness and the initial bud foraging aggressiveness.

For all multivariate models we used a Gaussian error structure for each response variable, 550,000 iterations, a burn in of 50,000, and a thinning interval of 100. Priors were set to be flat and relatively uninformative, with 70% of the phenotypic variance for the logged values of each trait placed on the residual variance, 20% on the among-colony variance, and 10% on the among-date variance (following: Brommer 2017).

## Results

### Robustness to disturbance

Across all treatments, pre-disturbance foraging aggressiveness showed consistent differences among colonies, (repeatability (r) of logged values = 0.152, credible intervals (CIs) = −0.060 to 0.348). Post-disturbance foraging aggressiveness was also consistently different among-colonies (r = 0.376, CIs = 0.158 to 0.555). We therefore conclude that each colony is in a relatively stable behavioural “state” of a particular level of foraging aggressiveness during the six days we measured them. The phenotypic correlation between pre- and post-disturbance foraging aggressiveness was significant and positive (r = 0.217, t = 2.321, df = 109, p = 0.022). At the among-colony level, pre-disturbance foraging aggressiveness positively covaried with post-disturbance foraging aggressiveness, although the 95% CIs of the among-colony covariance overlapped zero (covariance mode = 0.167, CIs = - 0.103 to 0.598, correlation mode = 0.547, CIs = −0.124 to 0.850). Full model results are provided in the supplementary materials (Table S1). These findings suggest that colony collective behaviour is stable over time.

The phenotypic correlation between pre- and post-disturbance foraging aggressiveness in the control group was quite strong and positive (Fig. 3a, r = 0.482, t = 3.483, df = 40, p = 0.001), absent in the procedural control group (Fig. 3b, r = −0.093, t = −0.567, df = 37, p = 0.574), and weakly positive but non-significant in the removal group (Fig. 3c, r = 0.151, t = 0.809, df = 28, p = 0.425). At the among-colony level, for the control group, there was a positive correlation between pre- and post-disturbance foraging aggressiveness (Fig. 3a, covariance mode = 0.245, CIs = −0.240 to 1.080, correlation mode = 0.701, CIs = −0.177 to 0.953), no correlation at all in the procedural control group (Fig. 3b, covariance mode = −0.001, CIs = −0.621 to 0.464, correlation mode = 0.051, CIs = - 0.773 to 0.726), and a weak positive correlation in the removal group (Fig. 3c, covariance mode = 0.094, CIs = −0.547 to 1.086, correlation mode = 0.637, CIs = =-0.595 to 0.925). Note that the CIs of all of these correlations overlap zero and hence each other. See Tables S2-4 in the supplementary materials for full model results. Among-colony correlations therefore largely matched the phenotypic correlations (Fig. 3a-c). These results collectively convey that perturbing colonies by removing individuals disrupted colony collective behaviour, especially if the individuals were subsequently returned.

**Figure 3.**
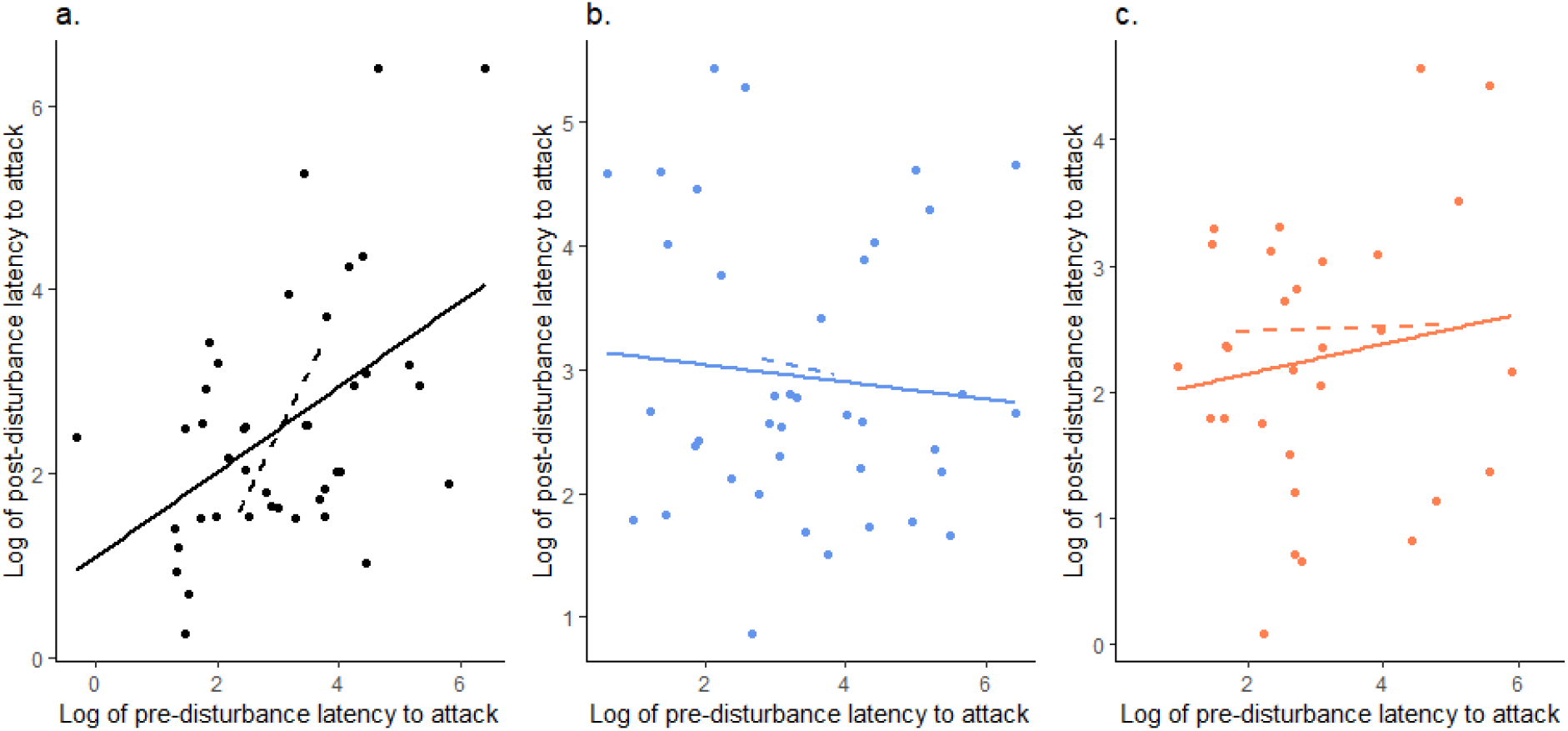
The relationship between the logs of pre- and post-disturbance foraging aggressiveness in the three treatment groups (a. control, b. procedural control, c. removal). Solid lines show the phenotypic correlations, while dashed lines showed the estimated among-colony correlations from the multivariate model.

### Transmission of collective behaviour

Colonies showed consistent differences in foraging aggressiveness in the laboratory (r = 0.282, CIs = 0.080 to 0.472). Bud colonies showed a small amount consistent differences in in the initial three measures of foraging aggressiveness (r = 0.082, CIs = 0.024 to 0.332). There was no phenotypic correlation between pre-disturbance foraging aggressiveness and initial bud foraging aggressiveness (Fig. 4a, r = 0.043, t = 0.272, df = 40, p = 0.787) or laboratory foraging aggressiveness (Fig. 4b, r = 0.065, t = 0.610, df = 88, p = 0.543). Laboratory and initial bud behaviour were also not correlated (Fig. 4c, r = −0.019, t = −0.120, df = 40, p = 0.905). Correlations were also absent at the among-colony level (pre-disturbance & initial bud foraging aggressiveness: Fig. 4a, covariance mode = 0.042, CIs = −0.314 to 0.502, correlation mode = 0.143, CIs = −0.553 to 0.814; pre-disturbance & laboratory foraging aggressiveness: Fig. 4b, covariance mode = 0.004, CIs = −0.342 to 0.431, correlation mode = 0.133, CIs = −0.504 to 0.651; laboratory & initial bud foraging aggressiveness: Fig. 4c, covariance mode = 0.008, CIs = −0.462 to 0.560, correlation mode = 0.386, CIs = −0.631 to 0.779). Full model results are given in the supplementary materials (Table S5).

**Figure 4.**
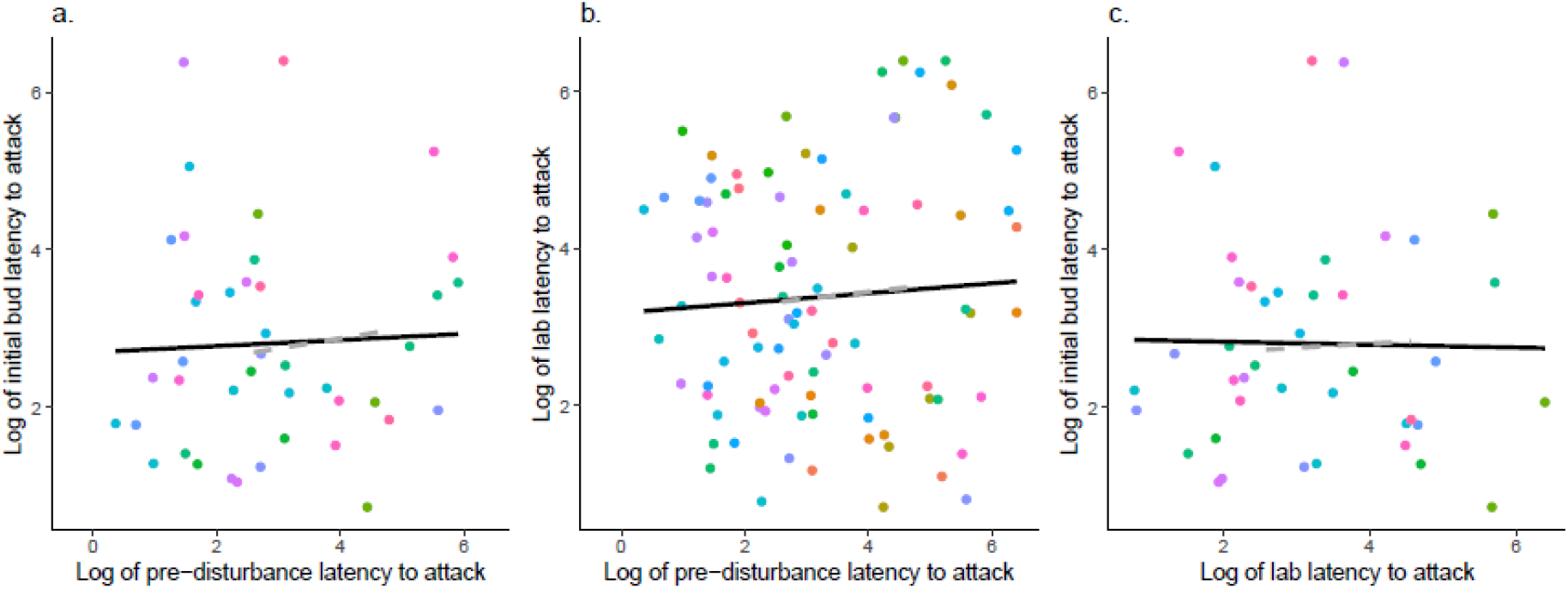
The relationship between a. pre-disturbance foraging aggressiveness and lab foraging aggressiveness, b. pre-disturbance foraging aggressiveness and initial bud foraging aggressiveness, and c. lab foraging aggressiveness and initial bud foraging aggressiveness. Point colours indicate different colonies. Solid black lines show the phenotypic correlations, while the dashed grey lines show the estimated among-colony correlations from the multivariate model.

Settled bud behaviour showed consistent differences among-colonies in foraging aggressiveness (r = 0.161, CIs = 0.044 to 0.464). There was a phenotypic correlation between settled bud behaviour and foraging aggressiveness (Fig. 5a, r = 0.464, t = 3.317, df = 40, p = 0.002), but not between settled bud behaviour and laboratory foraging aggressiveness (Fig. 5b, r = −0.117, t = −0.743, df = 40, p = 0.462). At the among-colony level, settled bud foraging aggressiveness was positively correlated with pre-disturbance foraging aggressiveness, although the CIs overlapped zero (Fig. 5a, covariance mode = 0.136, CIs = −0.214 to 0.696, correlation mode = 0.576, CIs = −0.269 to 0.896). Laboratory foraging aggressiveness was not correlated with settled bud foraging aggressiveness (Fig. 5b, covariance mode = 0.005, CIs = −0.534 to 0.549, correlation mode = 0.133, CIs = −0.675 to 0.736). Full model results are given in the supplementary materials (Table S6). Therefore, as for the robustness to disturbance, phenotypic correlations matched the among-colony correlations. These results suggest that parent and offspring colony collective behaviours can resemble each other, but only once the offspring colony had settled into an environment close to that of the parental colony’s.

**Figure 5.**
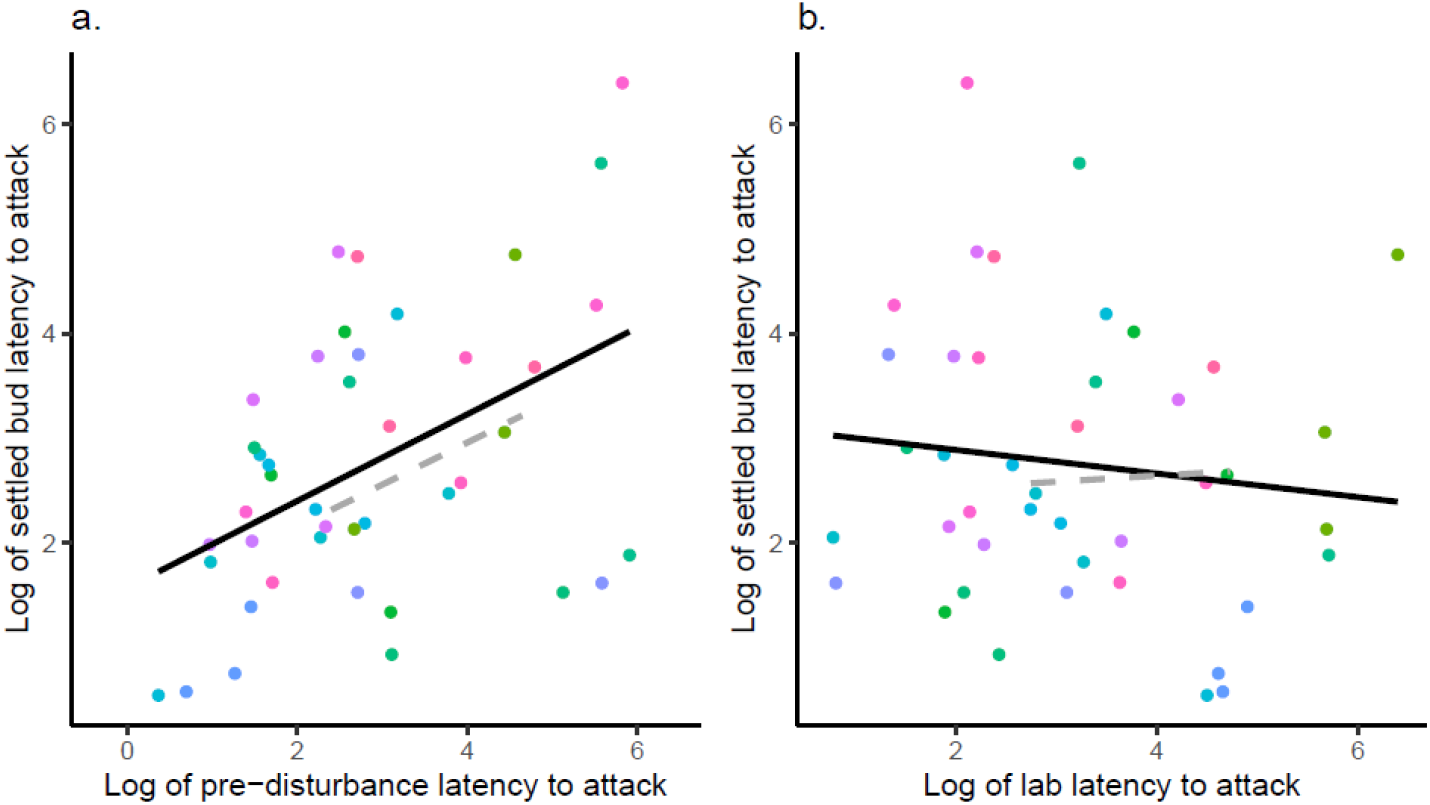
The relationship between a. pre-disturbance foraging aggressiveness and settled bud foraging aggressiveness, and b. lab foraging aggressiveness and settled bud foraging aggressiveness. Point colours indicate different colonies. Solid black lines show the phenotypic correlations, while the dashed grey lines show the estimated among-colony correlations.

The volume of the colony’s basket, number of adults, and trial number did not influence foraging aggressiveness in any of the models. There was some variation among days in foraging aggression, see Tables S1-6 for estimates.

## Discussion

Organisms in groups can possess collective behaviours, which can be subject to selection. How robust these collective behaviours are to disturbance, and whether they are transmitted from parent groups to offspring groups, is however not well known. Here we show that the foraging aggressiveness of *A. eximius* colonies is consistent over a period of several weeks and presumably longer, given that at high elevations foraging aggressiveness can influence colony survival over many months (Lichtenstein et al. 2019). Yet, colony behaviour is not robust to perturbation, especially if individuals are removed from the colony and then returned. We further found that bud colonies had collective behaviour that resembled that of their parent colony, but this was only apparent once the bud colony had spent over a week settling after the translocation and was not apparent when comparing laboratory measures of the bud colony with the parent colony.

First, we note here that, while the all patterns we detected in the study were qualitatively same at the among-colony level as at the phenotypic level, the 95% credible intervals of all among-colony correlations overlapped zero. From inspection of the correlation coefficients (see also Figs. 3–5), we can see the among-colony correlations are often stronger than the phenotypic correlations. Therefore, the overlap with zero is likely due to high uncertainty, probably due to our study using fewer than 50 colonies, and fewer than 20 colonies in each treatment group, rather than a small effect size. We therefore take the liberty of discussing among-colony correlations that are of the same strength or stronger than an equivalent and statistically significant phenotypic correlation. We do this because we consider these results to represent meaningful biological trends rather than statistical error.

### Collective behaviour is vulnerable to disturbance

There were consistent differences among colonies in both pre- and post-disturbance behaviour, but no covariance between pre- and post-disturbance behaviours in the procedural control and removal treatment groups. This suggests that foraging aggressiveness represents a semi-stable state that a colony is in, but that the colony is shifted to a different state by perturbations, as colonies did not retain the same level of foraging aggressiveness when individuals were removed or when the colony was disturbed by the removal and then return of individuals. Discussing populations or ecosystems as “systems” that can exist in different states has a long history in ecology (May 1974; Solé and Goodwin 2000). Referring to social groups in this way is less common, but interest in the utility of this viewpoint is growing (Flack et al. 2005, 2006; Doering et al. 2018; Pruitt et al. 2018). Social systems have previously been shown to be vulnerable to shifts from calm to antagonistic states due to the removal of key individuals (Flack et al. 2005, 2006) or due to gradual heating (Doering et al. 2018). Here we have found that the removal of individuals combined with a physical disturbance to the colony causes the colony to shift from one state of foraging aggression to another, although we did not observe a general increase in aggression due to the perturbations. In fact, mean foraging aggressiveness was equal in the control and removal treatment groups, and lower (longer latencies) in the procedural control group. We concluded this based on comparing the intercepts for post-disturbance foraging aggressiveness between the models for each treatment (although note that the 95% credible intervals overlapped in all cases, see Tables S2-4). Instead, we have observed that a colony adopts a different, yet still repeatable, behaviour to what it displayed before the disturbance.

As spider colonies did not return to their original foraging aggressiveness after the disturbance, consistent differences in behaviour among-colonies probably do not rely on some underlying stable trait of the colony (as is suggested for “pace of life syndrome” hypotheses for consistent among-individual differences in behaviour; (Réale et al. 2010)). Instead, consistent differences among colonies may depend on social interactions that generate positive feedback loops that cause colonies to diverge in behaviour (e.g. Luttbeg and Sih 2010). Such multiplicative interactions can give systems that are highly sensitive to initial conditions, and hence give variable trajectories and final states (Boyce 1992; Hastings et al. 1993; Cole 1994). Therefore, following the perturbation, *A. eximius* colonies may engage in interactions that, despite being deterministic and so giving rise to consistent behaviour, nevertheless follow divergent trajectories and so do not give the same behavioural trait as the colony previously possessed (Fisher et al. 2018). Interactions between individual *A. eximius* within the colony that catalyse increased aggression could give this dynamic, while interactions between the whole colony and its environment might also generate sufficient feedback. Currently, our understanding of the development of *A. eximius* colony collective behaviour is insufficient to allow us to judge the likely relative contributions of these two possibilities. However, social network analysis on the distance related social spider *Stegodypus dumicola* hint that positive feedback within colonies can cause the accentuation of individual differences within groups (Hunt et al. 2018), raising the possibility something similar could happen in *A. eximius*.

Removing individuals and then adding them back to the colony (as occurred in the procedural control group) completely removed any relationship between pre- and post-disturbance foraging aggressiveness. This suggests that removing individuals for a time and then returning them destabilises collective behaviour much more than simply removing them. The returning spiders may not have been recognised by their old colony-mates, and a period of antagonism may have disrupted colony behaviour. Social (and subsocial) spiders are thought to discriminate between kin and non-kin (Evans 1999; Bilde and Lubin 2001; Beavis et al. 2007; Schneider and Bilde 2008; Grinsted et al. 2011). However, *A. eximius* is known to accept intruders from the different colonies as well as from the same colony (Pasquet et al. 1997), suggesting there would have been limited antagonism towards the returning spiders. Instead, Pasquet *et al*. (1997) observed that the presence of an intruder increases the nearest neighbour distance within a colony. This change could then influence collective foraging aggressiveness. For now, we propose that the especially destabilised foraging behaviour of these colonies stems from their effectively experiencing two social disturbance as opposed to just one: having both lost a subset of group members and regained them, regardless of the familiarity of these group members.

### Parent and offspring colony collective behaviours resemble each other, but only once settled into the same environment

Parent colony behaviour (pre-disturbance) only covaried with bud colony behaviour once the bud colony had settled. This suggests that a group phenotype can be transmitted from parent to offspring colonies, like individual behaviours often are. However, this was only apparent over a week after the bud colony was been returned to the wild, suggesting there is an initial settling period before the bud colony regains the collective behaviour its parent colony showed. Further, parent colony foraging aggressiveness did not covary with laboratory foraging aggressiveness. Behaviour in the laboratory could therefore represent a different trait to behaviour in the wild, perhaps owing to colonies’ residing in completely different environments. In short, it could be that that bud colonies were permitted to reassume a shared environment that drives the correlation between parent colony and bud colony (Kruuk and Hadfield 2007). If this is so, then foraging aggressiveness might itself not be transmitted between parent and offspring groups.

To evaluate the possible influence of a shared environment, we need to identify an environmental variable that could drive such a parent-offspring resemblance (Kruuk and Hadfield 2007). Foraging aggressiveness in *A. eximius* decreases at higher elevations (Lichtenstein et al. 2019), and our study included colonies from 398m to 1146m above sea level (Fig.1). We tested whether it was elevation that drove the parent-offspring correlation by re-fitting the model for pre-disturbance and settled bud foraging aggressiveness (the model also contained laboratory foraging aggressiveness as a third response, but it is not important here) with the elevation of the colony (mean centred and scaled to a variance of one) as a fixed effect. In this model, pre-disturbance foraging aggression was lower (latencies tended to be longer) at higher elevations, although the credible intervals for the effect overlapped zero (fixed effect mode = 0.392, CIs = −0.059 to 0.836), but settled bud foraging aggressiveness did not change with elevation (fixed effect mode = 0.034, CIs = −1.001 to 1.261). In this model the relationship between pre-disturbance and settled bud foraging aggressiveness was roughly the same as in the model without elevation (covariance mode = 0.119, CIs = −0.212 to 0.715, correlation mode = 0.583, CIs = −0.283 to 0.892). This therefore suggests that sharing the same elevation was not driving the similarity between parent and offspring colonies. However, it is possible that other environmental variables are driving the resemblance.

An alternative explanation for the parent-offspring colony resemblance is that different colonies use different but repeatable behaviour rules to assemble colony behaviour. Therefore, once offspring colonies had settled, they were able to recreate the collective behaviour of the parent colony. While such a dynamic suggests a colony’s collective behaviour would resist a perturbation, it may take some time for the original collective behaviour to re-establish. Notably, extra time which was granted to the offspring colonies because of their conspicuous readjustments in the foliage, but not to the parent colonies post-disturbance. If we had tracked the parent colonies post-disturbance for a longer period of time, we may have seen their foraging aggressiveness return to its pre-disturbance level.

The outcome of selection on collective behaviour are quite different if collective behaviour is determined by an environmental variable (other than elevation) versus a directly transmitted quality of the parent colony. Relatedness within *A. eximius* colonies is typically very high (average r = 0.92 across four populations in Suriname, although r was estimated as 0.18 based on two nearby colonies at a site in Panama; (Smith and Hagen 1996)), and so selection at the colony level could be expected to give adaptation at the colony-level (Gardner and Grafen 2009; Queller and Strassmann 2009). If collective behaviour is determined by the environment, then selection will most likely favour colonies that best match their behaviour to the environment. In this case, changes to populations’ behaviour across generations is more likely to reflect changes in habitat availabilities or selection acting on some aspect of colonies’ habitat preferences or dispersal abilities. In contrast, if foraging aggressiveness is genuinely directly passed from parent colony to offspring colony, and given at high elevations we can observe selection against high foraging aggression (Lichtenstein et al. 2019), then we might expect mean aggression at high elevations to decrease across generations by selection acting directly on colony behaviour.

### Conclusions

In summary, we found that the foraging aggressiveness of *A. eximius* colonies is relatively stable over time but can be disrupted by perturbations. Returning individuals to their source colony disrupts a colony’s collective foraging even more than simply removing individuals from a colony. Offspring colonies have collective behaviour that resembles that of their parent colony, and this does not appear to be driven by a shared elevation. Instead, other forces like shared microhabitat preferences or the direct transmission of colony interaction rules, genetically determined behaviours, or plastic states (e.g., hunger levels, aggression levels) may drive resemblance of parent and offspring colonies. Appreciating that groups possess behavioural states, and that these states may be influenced by external perturbations yet still be passed from parent group to offspring group, should help us understand the role of group phenotypes in ecological and evolutionary processes.

## Author contributions

DNF, JLLL and JNP designed the study. JY acquired the permits. DNF, JLLL & RCP collected the data. DNF analysed the data and drafted the manuscript. All authors contributed to revisions of the manuscript and approved the final version.

## Acknowledgements

We thank J. B. Barnett, H. M. Anderson and B. L. McEwen for being excellent colleagues in the field. We also thank T. D. Swanson and the staff of the Andes and Amazon Field School at Iyarina for making our stay as comfortable and enjoyable as possible. Funding was provided by a Canada 150 Research Chair award to JNP. We have no conflicts of interest.

## Data accessibility

All data and R code used in the analysis will be made available upon publication.

