## Supplementary materials - Tables S1-6 for "Collective behaviour is not robust to disturbance, yet parent and offspring colonies resemble each other in social spiders"

| Variable | Posterior mode | Posterior mean | Lower 95% credible interval | Upper 95% credible interval | effect |
| --- | --- | --- | --- | --- | --- |
| Intercept: pre-disturbance | 3.264 | 3.194 | 2.587 | 3.810 | fixed |
| Intercept: post-disturbance | 2.626 | 2.591 | 2.159 | 3.062 | fixed |
| Trial: pre-disturbance | -0.141 | -0.164 | -0.828 | 0.535 | fixed |
| Trial: post-disturbance | -0.042 | -0.094 | -0.579 | 0.346 | fixed |
| Colony volume: pre-disturbance | 0.201 | 0.167 | -0.125 | 0.482 | fixed |
| Colony volume: post-disturbance | 0.119 | 0.108 | -0.196 | 0.417 | fixed |
| Among-colony variance: pre-disturbance | 0.373 | 0.501 | 0.121 | 0.915 | random |
| Among-colony covariance: pre- & post-disturbance | 0.167 | 0.227 | -0.103 | 0.598 | random |
| Among-colony variance: post-disturbance | 0.521 | 0.635 | 0.206 | 1.115 | random |
| Among-date variance: pre-disturbance | 0.173 | 0.430 | 0.030 | 1.102 | random |
| Among-colony variance: post-disturbance | 0.061 | 0.162 | 0.018 | 0.428 | random |
| Residual variance: post-disturbance | 1.645 | 1.629 | 1.191 | 2.121 | residual |
| Among-colony variance: pre-disturbance | 0.888 | 0.941 | 0.649 | 1.266 | residual |
| Pre- & post-disturbance correlation = 0.547, CIs = -0.124-0.850 | | | | | |

### Supplementary materials for “Collective behaviour is not robust to disturbance, yet parent and offspring colonies resemble each other in social spiders”

**Table S1**. Full model results for the covariance between pre- and post-disturbance behaviour across all treatments.

**Table S2**. Full model results for the covariance between pre- and post-disturbance behaviour for the control treatment.

| Variable | Posterior mode | Posterior mean | Lower 95% credible interval | Upper 95% credible interval | effect |
| --- | --- | --- | --- | --- | --- |
| Intercept: pre-disturbance | 2.992 | 3.021 | 2.279 | 3.797 | fixed |
| Intercept: post-disturbance | 2.430 | 2.516 | 1.788 | 3.172 | fixed |
| Trial: pre-disturbance | -0.322 | -0.323 | -1.023 | 0.483 | fixed |
| Trial: post-disturbance | 0.141 | 0.112 | -0.470 | 0.715 | fixed |
| Colony volume: pre-disturbance | 0.167 | 0.106 | -0.427 | 0.631 | fixed |
| Colony volume: post-disturbance | 0.466 | 0.496 | -0.055 | 1.084 | fixed |
| Among-colony variance: pre-disturbance | 0.390 | 0.587 | 0.091 | 1.391 | random |
| Among-colony covariance: pre- & post-disturbance | 0.245 | 0.358 | -0.240 | 1.080 | random |
| Among-colony variance: post-disturbance | 0.580 | 0.871 | 0.103 | 1.921 | random |
| Among-date variance: pre-disturbance | 0.160 | 0.412 | 0.032 | 1.193 | random |
| Among-colony variance: post-disturbance | 0.104 | 0.262 | 0.023 | 0.785 | random |
| Residual variance: pre-disturbance | 1.360 | 1.418 | 0.811 | 2.126 | residual |
| Residual variance: post-disturbance | 0.755 | 0.851 | 0.456 | 1.396 | residual |
| Pre- & post-disturbance correlation = 0.701, CIs = -0.177 – 0.953 | | | | | |

**Table S3**. Full model results for the covariance between pre- and post-disturbance behaviour for the procedural control treatment.

| Variable | Posterior mode | Posterior mean | Lower 95% credible interval | Upper 95% credible interval | effect |
| --- | --- | --- | --- | --- | --- |
| Intercept: pre-disturbance | 3.358 | 3.365 | 2.384 | 4.308 | fixed |
| Intercept: post-disturbance | 3.003 | 2.945 | 2.343 | 3.567 | fixed |
| Trial: pre-disturbance | -0.129 | -0.150 | -1.166 | 0.907 | fixed |
| Trial: post-disturbance | -0.237 | -0.280 | -0.837 | 0.311 | fixed |
| Colony volume: pre-disturbance | 0.190 | 0.220 | -0.350 | 0.811 | fixed |
| Colony volume: post-disturbance | 0.108 | 0.086 | -0.411 | 0.594 | fixed |
| Among-colony variance: pre-disturbance | 0.271 | 0.609 | 0.083 | 1.456 | random |
| Among-colony covariance: pre- & post-disturbance | -0.000 | -0.029 | -0.621 | 0.464 | random |
| Among-colony variance: post-disturbance | 0.222 | 0.460 | 0.052 | 1.094 | random |
| Among-date variance: pre-disturbance | 0.269 | 0.855 | 0.028 | 2.398 | random |
| Among-colony variance: post-disturbance | 0.0813 | 0.184 | 0.021 | 0.505 | random |
| Residual variance: pre-disturbance | 1.240 | 1.549 | 0.776 | 2.445 | residual |
| Residual variance: post-disturbance | 0.934 | 1.128 | 0.582 | 1.752 | residual |
| Pre- & post-disturbance correlation = 0.051, CIs = -0.773 – 0.726 | | | | | |

**Table S4**. Full model results for the covariance between pre- and post-disturbance behaviour for the removal treatment.

| Variable | Posterior mode | Posterior mean | Lower 95% credible interval | Upper 95% credible interval | effect |
| --- | --- | --- | --- | --- | --- |
| Intercept: pre-disturbance | 2.973 | 3.017 | 2.240 | 3.850 | fixed |
| Intercept: post-disturbance | 2.500 | 2.390 | 1.662 | 3.160 | fixed |
| Trial: pre-disturbance | 2.73 | 0.144 | -0.647 | 0.895 | fixed |
| Trial: post-disturbance | -0.164 | -0.140 | -0.865 | 0.524 | fixed |
| Colony volume: pre-disturbance | 0.262 | 0.236 | -0.410 | 0.818 | fixed |
| Colony volume: post-disturbance | -0.164 | -0.268 | -0.892 | 0.318 | fixed |
| Among-colony variance: pre-disturbance | 0.567 | 1.011 | 0.126 | 2.195 | random |
| Among-colony covariance: pre- & post-disturbance | 0.094 | 0.194 | -0.547 | 1.086 | random |
| Among-colony variance: post-disturbance | 0.196 | 0.501 | 0.051 | 1.301 | random |
| Among-date variance: pre-disturbance | 0.111 | 0.298 | 0.026 | 0.861 | random |
| Among-colony variance: post-disturbance | 0.085 | 0.265 | 0.019 | 0.803 | random |
| Residual variance: pre-disturbance | 1.643 | 1.820 | 1.020 | 2.749 | residual |
| Residual variance: post-disturbance | 0.0805 | 0.977 | 0.443 | 1.604 | residual |
| Pre- & post-disturbance correlation = 0.637, CIs = -0.595 – 0.925 | | | | | |

**Table S5**. Full model results for the covariance between pre-disturbance, laboratory, and initial bud behaviour

| Variable | Posterior mode | Posterior mean | Lower 95% credible interval | Upper 95% credible interval | | effect |
| --- | --- | --- | --- | --- | --- | --- |
| Intercept: pre-disturbance | 3.202 | 3.199 | 2.606 | | 3.838 | fixed |
| Intercept: laboratory | 3.385 | 3.521 | 2.959 | | 4.138 | fixed |
| Intercept: initial bud | 2.788 | 2.754 | 1.855 | | 3.646 | fixed |
| Trial: pre-disturbance | -0.105 | -0.165 | -0.859 | | 0.572 | fixed |
| Trial: laboratory | -0.452 | -0.386 | -0.936 | | 0.168 | fixed |
| Trial: initial bud | -0.001 | 0.015 | -0.871 | | 1.020 | fixed |
| Colony volume: pre-disturbance | 0.144 | 0.165 | -0.121 | | 0.469 | fixed |
| Number of adults: laboratory | -0.234 | -0.253 | -0.701 | | 0.197 | fixed |
| Number of adults: initial bud | -0.072 | -0.089 | -0.910 | | 0.763 | fixed |
| Among-colony variance: pre-disturbance | 0.441 | 0.506 | 0.138 | | 0.954 | random |
| Among-colony covariance: pre- disturbance & laboratory | 0.004 | 0.034 | -0.342 | | 0.431 | random |
| Among-colony covariance: pre-disturbance & initial bud | 0.042 | 0.078 | -0.314 | | 0.502 | random |
| Among-colony variance: laboratory | 0.611 | 0.743 | 0.145 | | 1.461 | random |
| Among-colony covariance: laboratory & initial bud | 0.008 | 0.056 | -0.462 | | 0.560 | random |
| Among-colony variance: initial bud | 0.246 | 0.416 | 0.059 | | 0.984 | random |
| Among-date variance: pre-disturbance | 0.209 | 0.448 | 0.031 | | 1.162 | random |
| Among-date variance: laboratory | 0.110 | 0.200 | 0.026 | | 0.510 | random |
| Among-date variance: initial bud | 0.129 | 0.499 | 0.026 | | 1.610 | random |
| Residual variance: pre-disturbance | 1.573 | 1.636 | 1.176 | | 2.104 | residual |
| Residual variance: laboratory | 1.607 | 1.662 | 1.096 | | 2.297 | residual |
| Residual variance: initial bud | 1.468 | 1.751 | 0.974 | | 2.660 | residual |
| Pre-disturbance & laboratory correlation = 0.133, CIs = -0.504 – 0.651 | | | | | | |
| Pre-disturbance & initial bud correlation = 0.143, CIs = -0.553 – 0.814 | | | | | | |
| Laboratory & initial bud correlation = 0.386, CIs = -0.631 – 0.779 | | | | | | |

**Table S6**. Full model results for the covariance between pre-disturbance, laboratory, and settled bud behaviour.

| Variable | Posterior mode | Posterior mean | Lower 95% credible interval | Upper 95% credible interval | effect |
| --- | --- | --- | --- | --- | --- |
| Intercept: pre-disturbance | 3.117 | 3.202 | 2.621 | 3.819 | fixed |
| Intercept: laboratory | 3.525 | 3.518 | 2.917 | 4.084 | fixed |
| Intercept: settled bud | 2.625 | 2.719 | 1.924 | 3.520 | fixed |
| Trial: pre-disturbance | -0.147 | -0.154 | -0.853 | 0.581 | fixed |
| Trial: laboratory | -0.334 | -0.376 | -0.951 | 0.194 | fixed |
| Trial: settled bud | 0.211 | 0.275 | -0.509 | 0.996 | fixed |
| Colony volume: pre-disturbance | 0.188 | 0.162 | -0.131 | 0.461 | fixed |
| Number of adults: laboratory | -0.338 | -0.243 | -0.662 | 0.233 | fixed |
| Number of adults: settled bud | 0.441 | 0.300 | -0.578 | 1.173 | fixed |
| Among-colony variance: pre-disturbance | 0.467 | 0.532 | 0.122 | 0.967 | random |
| Among-colony covariance: pre- disturbance & laboratory | 0.090 | 0.023 | -0.354 | 0.427 | random |
| Among-colony covariance: pre-disturbance & settled bud | 0.136 | 0.216 | -0.214 | 0.696 | random |
| Among-colony variance: laboratory | 0.550 | 0.730 | 0.176 | 1.432 | random |
| Among-colony covariance: laboratory & settled bud | 0.005 | 0.030 | -0.534 | 0.549 | random |
| Among-colony variance: settled bud | 0.273 | 0.564 | 0.071 | 1.258 | random |
| Among-date variance: pre-disturbance | 0.189 | 0.447 | 0.042 | 1.167 | random |
| Among-date variance: laboratory | 0.081 | 0.204 | 0.028 | 0.517 | random |
| Among-date variance: settled bud | 0.088 | 0.290 | 0.023 | 0.885 | random |
| Residual variance: pre-disturbance | 1.529 | 1.615 | 1.170 | 2.097 | residual |
| Residual variance: laboratory | 1.556 | 1.662 | 1.107 | 2.294 | residual |
| Residual variance: settled bud | 1.321 | 1.504 | 0.840 | 2.316 | residual |
| Pre-disturbance laboratory correlation = -0.066, CIs = -0.534 – 0.617 | | | | | |
| Pre-disturbance settled bud correlation = 0.576, CIs = -0.269 – 0.896 | | | | | |
| Laboratory settled bud correlation = 0.133, CIs = -0.675 – 0.736 | | | | | |
